# Viral methyltransferase inhibitors: berbamine, venetoclax, and ponatinib as efficacious antivirals against Chikungunya virus

**DOI:** 10.1101/2024.05.19.594723

**Authors:** Mandar Bhutkar, Ankita Saha, Shailly Tomar

## Abstract

Chikungunya virus (CHIKV), transmitted by mosquitoes, poses a significant global health threat. Presently, no effective treatment options are available to reduce the disease burden. The lack of approved therapeutics against CHIKV and the complex spectrum of chronic musculoskeletal and neurological manifestations raise significant concerns, and repurposing drugs could offer swift avenues in the development of effective treatment strategies. RNA capping is a crucial step meditated by non-structural protein 1 (nsP1) in CHIKV replication. In this study, FDA-approved antivirals targeting CHIKV nsP1 methyltransferase (MTase) have been identified by structure-based virtual screening. Berbamine Hydrochloride (BH), ABT199/Venetoclax (ABT), and Ponatinib (PT) were the top hits, which exhibited robust binding energies. Tryptophan fluorescence spectroscopy-based assay confirmed binding of BH-, ABT-, and PT to purified nsP1 with K_D_ values ∼5.45 μM, ∼161.3 μM, and ∼3.83μM, respectively. A dose-dependent decrease in CHIKV nsP1 MTase activity was observed in a capillary electrophoresis-based assay. Treatment with BH, ABT, and PT lead to a dose-dependent reduction in the virus titer with IC_50_ <100, ∼3.46, and <3.9 nM, respectively, and reduced viral mRNA levels. The nsP1 MTases are highly conserved among alphaviruses; therefore, BH, ABT, and PT, as expected, inhibited replication machinery in Sindbis virus (SINV) replicon assay with IC_50_ ∼1.94, ∼0.23, and >1.25 μM, respectively. These results underscore the efficacy and promise of repurposing drugs as rapid and effective antiviral therapeutics against CHIKV.

**Highlights:**

- Berbamine Hydrochloride (BH), ABT199/Venetoclax (ABT), and Ponatinib (PT) identified as inhibitors of Chikungunya virus (CHIKV) nsP1
- Molecules BH, ABT and PT make moelcular interactions with nsP1: K_D_ values ∼5.45, ∼161.3, and ∼3.83 in μM, respectively and effectively inhibit methyltransferase activity.
- BH, ABT, and PT exhibited potent antiviral activity against CHIKV in cell-based assays, with IC_50_ <100, ∼3.46, and <3.9 nM.
- BH, ABT, and PT also inhibit Sindbis virus, the prototype alphavirus with IC_50_ ∼1.94, ∼0.23, and >1.25 μM respectively demonstrating broad spectrum antiviral efficacy.

**Graphical Abstract:** 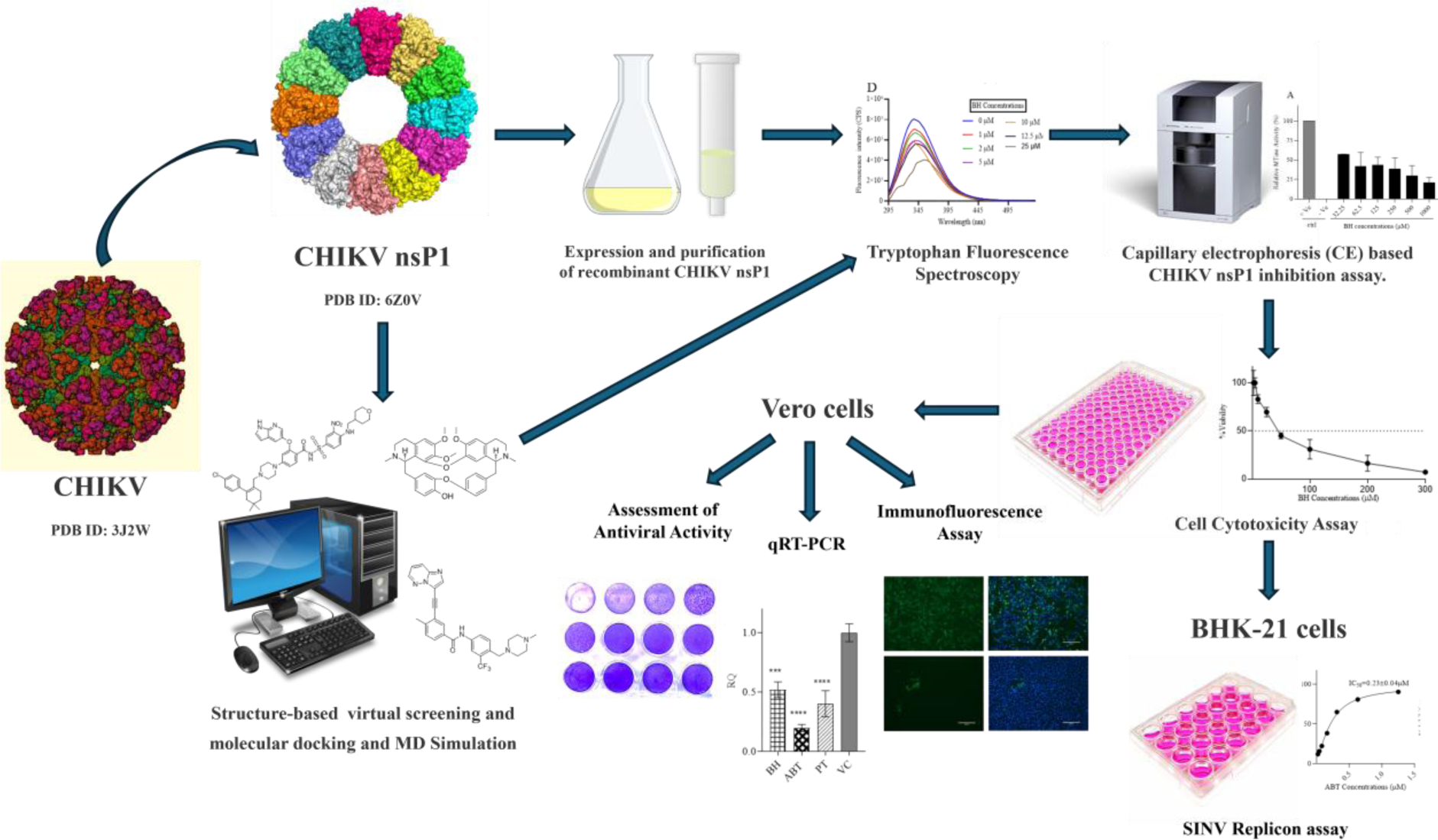

## Introduction

The *Togaviridae* family poses a formidable threat to human health. It comprises various viruses, such as Chikungunya virus (CHIKV), Sindbis virus (SINV), Semliki Forest virus (SFV), Rubella virus, etc (1). CHIKV belongs to the genus *Alphavirus* and is transmitted by *Aedes* mosquitos. It is an enveloped virus with a ∼12 kb positive-sense single-stranded RNA genome (+ssRNA) (2). CHIKV infection is associated with neurological complications and musculoskeletal manifestations characterized by fever, arthralgia, joint pain, and rash. CHIKV is responsible for various sporadic and local outbreaks worldwide (3). In 2023, the global burden of CHIKV disease reached alarming levels, with over half a million reported cases and a substantial number of fatalities worldwide (4,5). Furthermore, in 2022, ∼ 0.15 million CHIKV cases were reported in India (6). Although the FDA has approved a vaccine to prevent disease caused by CHIKV, ongoing concerns persist regarding insufficient demographic data, a 1.5% incidence of severe adverse reactions, and uncertainties surrounding its efficacy against chronic disease. Additionally, the vaccine is restricted to individuals aged 18 and older, limiting its applicability (7). Hence, there is an imminent need for effective antiviral therapy to treat CHIKV infection.

The alphaviruses replicate in the cytoplasm and have two open reading frames (ORF). The first ORF encodes for four non-structural proteins (nsP1, nsP2, nsP3, and nsP4), whereas the second ORF encodes structural proteins (capsid [C], envelope [E1, E2, E3], and transmembrane protein [6K]) (8). Of these, the nsP1 has N7-guanine methyltransferase (MTase) and guanylyl transferase (GTase) activities, which play an essential role in the life cycle of the virus. The nsP2 is a viral protease that post-translationally cleaves the viral polyprotein precursor. Moreover, it also possesses an N-terminal RNA triphosphatase/helicase activity, which aids in the unwinding of double-stranded RNA (9,10). Together with nsP2, the nsP1 protein caps viral RNA transcripts, protecting them from cellular exonucleases. In addition to this, capping facilitates viral mRNA recognition by eukaryotic translational initiation factor 4E (eIF4E) to initiate translation (11). Furthermore, the cap also aids the viral genome to evade recognition from the host innate immune system pathways such as retinoic acid-inducible gene I (RIG-I) and melanoma differentiation-associated protein 5 (MDA5) (12). Recently, nsP1’s role in decapping the cellular RNA has been demonstrated, facilitating its susceptibility to cellular exonucleases, and regulating cellular mRNA gene expression. In decapping, nsP1 has a mechanism analogous to the influenza virus ‘cap snatching’ mechanism (13).

In eukaryotic cells, cellular enzymes use conventional capping mechanisms, where guanosine triphosphate (GTP) is first transferred to the 5’ end of RNA, followed by its methylation. Contrastingly, in the case of alphaviruses, the capping of viral RNAs is an unconventional mechanism (14). In alphaviruses, viral RNA capping occurs in two significant steps. First, S-adenosyl methionine (SAM) acts as a methyl donor, transferring a methyl group to the seventh position of GTP. The methylation steps lead to the formation of m7 guanosine-5’-monophosphate (m7GMP), where S-adenosyl homocysteine (SAH) is generated as a by-product. This is known as the methyltransferase (MTase) step of the capping reaction (8). Following the methylation step, the m7GMP is covalently attached to the nsP1 enzyme via a single bond with His37 during the second step, known as guanylation (GT) step (15). Interestingly, this two-step mechanism yields a distinct cap 0 structure on alphaviral RNA characterized by monomethylation at the N7 position of the guanosine nucleotide (8).

Recently, the CHIKV nsP1 dodecameric Cryo-EM structure (6Z0V, 6Z0U, and 7DOP) has been published (16,17). This structure has an upper ring with MTase/GTase bifunctional catalytic activity. The lower ring has a role in oligomerization and membrane binding (16). Hence, this structure has increased the utility of high-throughput computational screening of small molecules for identifying novel drug candidates (16,17). Identifying novel antiviral compounds is an intricate, resource and time-intensive process. Drug repurposing involves identifying new uses for approved or investigational drugs beyond their original medical indications (18). This strategy takes advantage of completed early-stage trials, reducing the risks of failure and shortening the timeframe and financial investment required for drug development. Hence, structure-based virtual screening is a promising approach to identify compounds with potential pharmacological activity.

The present study profiled a library of drugs encompassing ∼3180 FDA small molecules, utilizing a combination of virtual screening methodologies and experimental validation approaches to identify molecules with potent inhibitory activities against CHIKV. Using this approach, Berbamine hydrochloride (BH), ABT-199/ Venetoclax (ABT), and Ponatinib (PT) were identified as top hits exhibiting dose-dependent antiviral activity against CHIKV. Assessing the *in vivo* efficacy of these molecules in clinical settings could present an important opportunity to fast-track the development of potential remedies against Alphaviruses.

## Materials and methods

### Multiple Sequence Alignment (MSA)

The amino acid sequence of nsP1 protein of alphaviruses was compared with CHIKV nsP1 as a reference point using Clustal Omega(19) to check if the key residues involved in the capping of viral RNA were conserved across different viruses. The MSA was performed for Mayaro virus (MV) (AZM66145.1), SFV (NC_003215.1), Barmah forest virus (BFV) (QVM79755.1), Eilat virus (EV) (NC_018615), CHIKV (QOW97289.1), Madariaga virus (MDV) (ABL84688.1), Aura virus (AV) (AWQ38330.1), Salmonid alphavirus (SAV) (KC122922.1), Ross River virus (RRV) (QTC33397.1), Venezuelan equine encephalitis virus (VEEV) (AAU89533.1), SINV (AWT57896.1), and Middelburg virus (MBV) (QOY44469.1) from the *Alphavirus* family. The sequence alignment profile of the selected nsP1 sequences was performed via Clustal Omega tool and analyzed by a graphical coloured depiction using ESPript 3.0 (20).

### Structure-based virtual screening and molecular docking

The cryo-electron microscopy (cryo-EM) structure of CHIKV nsP1 (6Z0V, 6Z0U, and 7DOP) revealed the presence of SAM and GTP binding catalytic site in nsP1 (17). Utilizing the 3-dimensional structure as a template, the present study employed PyRx 0.8 virtual screening tool to screen FDA-approved compound library centering around the active site of CHIKV nsP1 (21). For this purpose, the structure of CHIKV nsP1 was modelled using SWISS-MODEL since the atomic coordinates for several flexible loops were missing in the original PDB entry (6Z0U). The sdf files of compounds were virtually screened against the catalytic pocket. The grid box dimensions used for this screening were 33.48 Å × 19.99 Å × 32.10 Å, centered at coordinates X = 54.92, Y = 137.80, and Z = 91.35. Top-hit compounds targeting specific CHIKV nsP1 were selected for further redocking studies by AutoDock Vina (22). Briefly, the structures of ligands, i.e., BH, ABT, and PT, were obtained from the PubChem site in .sdf format and then converted to. pdbqt format using OpenBabel in PyRx 0.8 algorithm (21,23). In AutoDock Vina, grid dimensions were set to match those previously specified. The conformations with the best binding energy were further analyzed in PyMol and LigPlot+ software (24,25). Molecules demonstrating strong binding affinities and significant interactions with target pockets were prioritized for in-depth interaction analysis and experimental validation.

### Molecular Dynamics (MD) Simulation

Protein-ligand complexes of nsP1 with three top-hits, BH, ABT, and PT, identified from virtual screening studies were subjected to MD simulation studies to assess the flexibility and stability of protein-ligand interactions. GROMACS 2022.2 suite was used to conduct all simulation studies using the Charmm8 force field on a LINUX-based workstation (26). Ligand parameters and topology files were generated using the CHARMM General Force Field (CgenFF) program (27). Ions and water molecules were added to neutralize the whole cubic system for solvation. The energy minimization step was performed using the steepest descent method, followed by a two-phased equilibration of a constant number of particles, volume, and temperature (NVT), a constant number of particles, pressure, and temperature (NPT). NVT equilibration was done at 300K with a short-range electrostatic cut-off of 1.2 nm, and temperature regulation was done using the Berendsen temperature coupling method. This was followed by the next phase of equilibration NPT, and coordinates were generated every 1 ps. Finally, a 100 ns MD production run was performed with an integration time frame of 2fs, and the trajectories were generated after every ten ps. The conformations generated during the production step were used for calculating the root mean square deviation (RMSD) of protein-ligand complexes.

### Preparation of BH, ABT, and PT stock solutions

BH was purchased from Chemfaces (China) while ABT and PT were purchased from BLD Pharma (India). For experiments, the stock solutions (BH: 146.7 mM ABT: 57.57 mM; PT: 144.44 mM) for all compounds were prepared in dimethyl sulfoxide (DMSO) (Sigma-Aldrich) and filtered through a 0.2 μm size syringe filter (Millipore). Further dilutions were prepared in experimental buffers at the time of usage.

### Expression and purification of recombinant CHIKV nsP1

A single colony of *E. coli* Rosetta cells transformed with the recombinant expression plasmid pET28c-nsP1CHIKV was inoculated in a tube containing 10 mL of LB medium supplemented with 50 μg/mL kanamycin, 35 μg/mL of chloramphenicol, and cultured overnight at 37 °C and 180 rpm in a shaking incubator. The primary culture was used as inoculum for a secondary culture of 1L of the same medium. The flask was incubated at 37 °C and 180 rpm until the OD_600_ of the culture reached 0.4. The temperature was reduced at this point to 18 °C. Growth was continued at 18 °C until the optical density of 0.7 at OD_600_ was reached. 0.4 mM isopropyl-ß-thiogalactopyranoside (IPTG) was used to induce protein expression. After induction with IPTG, the culture was further grown for 12 h at 18 °C. Cells were harvested by centrifugation at 5000g at 4 °C. The obtained cell pellets were resuspended in 30 mL of binding buffer (50 mM Tris−HCl pH 7.3, 20 mM imidazole, 5% glycerol, and 200 mM NaCl), followed by cell lysis using French Press (Constant Systems, Ltd., Daventry, England). The obtained cell lysate was clarified by centrifugation at 12,000g for 45 min at 4 °C. Ni-nitrilotriacetic acid (NTA) beads (Qiagen, USA) were used to purify the protein. The supernatant was applied onto the pre-equilibrated Ni-NTA column and incubated for 30 min at 4 °C. After incubation, the column was washed using a buffer containing a gradient of increasing imidazole concentrations. Elution of the recombinant nsP1 protein was done in 50 mM Tris-HCl pH 7.3, 250 mM imidazole, 5% glycerol, 200 mM NaCl, and the fractions were analyzed by Sodium dodecyl-sulfate polyacrylamide gel electrophoresis (SDS-PAGE) and Coomassie blue staining. The purified protein fractions were pooled and dialyzed against the dialysis buffer (50 mM Tris-HCl pH 7.3, 5% glycerol, 100 mM NaCl) at 4 °C for 3 h. Further, CHIKV nsP1 protein was concentrated to the required concentration with Amicon centrifugal filters (10,000 MWCO Millipore, Burlington, MA, USA), flash-frozen in liquid nitrogen, and stored at −80 °C (15,28,29).

### Tryptophan Fluorescence Spectroscopy (TFS)

TFS experiments were conducted using a Fluoromax fluorescence spectrophotometer (Horiba Scientific). A quartz cuvette with dimensions of 5×5 mm was utilized. The excitation wavelength was set to 280 nm, and the emission wavelength was scanned from 295 to 540 nm. A slit width of 5 nm was employed for all measurements. Recombinant CHIKV nsP1 protein samples were prepared at a concentration of 1 μM in a 500 μL phosphate-buffered saline (PBS) solution. BH, ABT, and PT were titrated into the protein solution to achieve concentration ranges of 0 to 25 µM, 0 to 300 µM, and 0 to 5 µM, respectively. The experiments were conducted at a constant temperature of 25 °C. Control buffer experiments and compound titrations were performed parallel with the main experiments for background determination. Data from three independent experiments were collected and analyzed using nonlinear regression with the ‘One Site-Specific Binding’ model. The data was analyzed using GraphPad Prism 8 software (30).

### Capillary electrophoresis (CE) based CHIKV nsP1 inhibition assay

In the investigation of CHIKV nsP1 inhibition by BH, ABT, and PT enzyme reactions were conducted using specific reaction mixtures. The reaction mixture for nsP1 consisted of 50 mM Tris buffer (pH 7.5), 10 mM KCl, 2 mM DTT, 2 mM MgCl_2_, 0.3 mM SAM, and 0.3 mM GTP, along with 5 μM nsP1 protein. These reactions were performed at 37 °C for 1 h. To establish negative controls, enzyme assays included a reaction with no SAM-GTP (substrate). After the incubation reaction was stopped by adding acetonitrile in 1:2 (vol/vol), 1 mM caffeine was used as an internal control (15,29). The mixture was vortex-mixed for 15 s, and the protein was precipitated for 20 min at 18,500 g. The supernatant was transferred to sample vials for CE analysis, as mentioned in Mudgal et al. (2020). All reactions were performed in triplicates (15,29).

### Cell line and Virus propagation

BHK-21 and Vero cell lines were obtained from NCCS, Pune, India. Cells were maintained in Dulbecco’s Modified Eagle Medium (DMEM; HiMedia) supplemented with 10% heat-inactivated fetal bovine serum (FBS; Gibco, Waltham, MA, USA) and 1X penicillin-streptomycin solution (HiMedia). A reporter Sindbis Virus (SINV) replicon (SINV-REP) was constructed in which all the structural proteins of SINV downstream of subgenomic (SG) promoter were replaced with Firefly luciferase gene (FLuc) and internal ribosome entry site element (IRES) element at the 3’ end. Vero cells were used for the propagation and titration of CHIKV. CHIKV (Accession No. KY057363.1) was propagated and titrated using the protocol reported by Singh et al., 2018 and then stored at −80 °C for further experiments (31).

### Cell Cytotoxicity Assay

Different concentrations of BH, ABT, and PT were evaluated for cytotoxicity on Vero cells using 3-(4,5-dimethyl-thiazol-2-yl)-2,5-diphenyltetrazolium bromide (MTT) (Himedia) assay. Vero cells were seeded in a 96-well plate and incubated till a confluency of 90% was attained. The cells were treated with different dilutions of the compound and incubated at 37 °C in a 5% CO_2_ incubator for 24 h. Similarly, BHK-21 cells were treated with varying compound concentrations for 24 h. Post incubation, 20 μL/well of MTT (5 mg/mL) was added per well and incubated for 4 h at 37 °C in a 5% CO_2_ incubator. After incubation, 150 μL/well of DMSO was added to dissolve formazan crystals. Plates were read at a wavelength of 570 nm using a Synergy BioTek multi-mode plate reader (BioTek Instruments, Inc.). The average absorbance of 0.1 % DMSO-treated cells was used as cell control. Data from three independent experiments were collected and analyzed (30). The 50% cytotoxic concentration (CC_50_) was determined based on a linear dose-response analysis using GraphPad Prism 8 software.

### Assessment of Antiviral Activity

Vero cells were seeded in a 24-well plate at a cell density of 1.0 x 10^5^ cells/well. Cells were infected with CHIKV at a multiplicity of infection (MOI) of 0.1 and further incubated at 37 °C and 5% CO_2_ with gentle shaking every 15 min for 2 h. The inoculum was removed, and the cell monolayer was washed twice to ensure no chance of secondary infection. Compounds were prepared with 2% DMEM media, added post-infection (pi), and incubated at 37 °C and 5% CO_2_ for 24 h. Post incubation, the supernatant was collected and stored at -80 ^°^C, and plaque assay was performed as reported previously (31).

### Quantitative Real-Time PCR (qRT-PCR)

The treatment and infection procedures for BH, ABT, and PT compounds were conducted as mentioned in the assessment of antiviral activity. After the termination of the assay, trizol (Takara Bio) was added to the plate. RNA was purified according to the manufacturer’s protocol. Purified RNA was quantified by Nanodrop One Microvolume UV-Vis Spectrophotometer (Thermo Scientific). The quantified 400 ng of RNA used for cDNA preparation using the PrimeScript 1st strand cDNA Synthesis Kit (Takara Bio). The forward and reverse primers used for amplification are used as previously described (32). β-actin was utilised as an internal control. qRT-PCR was performed using KAPA SYBR fast universal qPCR kit on QuantStudio™ 5 System (Applied Biosystems, Waltham, MA, USA). The reactions were performed in triplicate, and the viral RNA load was assessed using the ΔΔCt method to derive the relative quantification (RQ) value (RQ = 2^(-ΔΔCt)). The decrease in viral RNA copy numbers following treatment with the three compounds was compared to an untreated virus control (32,33).

### Immunofluorescence Assay

Before a day of the infection, Vero cells were seeded in a 6-well plate at a cell density of 1 x 10^6^ cells/ well. Cells were treated with compounds, as mentioned earlier. The cells were washed three times with PBS and then fixed with methanol: acetone (1:1) for 1 h at room temperature, followed by permeabilization with 0.1% Triton-X-100. After washing, cells were incubated with antibodies against CHIKV (anti-alphavirus 1:100, SANTA Cruz Biotechnology Inc.) for 1 h and then incubated with fluorescein (FITC)-conjugated secondary anti-mouse antibody (1:250, Sigma) for 30 min at 37 °C. The plate was then rinsed with PBS and counter-stained with 4′,6-diamidino-2-phenylindole DAPI (Sigma) for 15 min. The images were captured under a fluorescence microscope (EVOS, Thermofisher) (30).

### SINV Replicon assay

Sindbis Virus Replicon (SINV-REP) assay was performed as described previously (29). SINV-REP construct derived from pToto64 clone encoding SINV non-structural proteins and a reporter luciferase tag was linearized using SacI (New England Biolabs), followed by *in vitro* transcription using SP6 RNA polymerase (New England Biolabs). 10 µg Capped mRNA transcripts were electroporated into BHK-21 cells in a 0.2 cm cuvette (25 μF, 1.5 kV, and 200 Ω) with the help of Gene Pulser Xcell Electroporation system (Bio-Rad) and resuspended in antibiotic-free DMEM supplemented with 10% FBS. Following this, the cells were seeded in a 24-well plate and incubated at 37 °C and 5% CO_2_ for 4 h. Post-incubation, the compound dilutions prepared in DMEM supplemented with 2% FBS were added to the cells and the cells were incubated for 6 h. The cells were lysed with lysis reagent (Promega) and the cell lysate was collected. The luciferase activity of cell lysates was measured using a firefly luciferase assay kit (Promega, USA), according to manufacturer’s instructions. Luminescence reading was measured using a Synergy BioTek multi-mode plate reader (BioTek Instruments, Inc.), 0.1% DMSO-treated cells were included as positive control, and untreated cells were included as cell control. The [inhibitor] v/s response curve was plotted in GraphPad Prism software.

### Statistical analysis

GraphPad Prism 8 software is used for data analysis. One-way ANOVA was used to determine statistical significance, followed by Dunnett’s tests wherever mentioned.

## Results

### Structure-guided identification of MTase inhibitors

The availability of cryo-EM atomic structure of CHIKV nsP1 capping enzyme facilitated *in silico* structure-based identification and characterization of BH, ABT, and PT as potential nsP1 inhibitors (16,34). In nsP1, the SAM binding site in nsP1 is lined by Gly65, Ser66, Ala67, Pro83, Arg85, Ser86, Asp89, Thr137, Asp138, and Gln151 residues. The GTP binding site comprises residues Ala40, Arg41, Ser44, Tyr154, Phe178, Phe241, Val243, Thr246, Tyr248, and Glu250. Arg70, Arg92, and Asp152 are common residues to both SAM and GTP binding site (16). MV, SFV, BFV, CHIKV, MDV, AV, RRV, VEEV, SINV, and MBV are known alphaviruses that cause human infection (19). In contrast, SAV exhibits tropism for Atlantic salmon, causing pancreatic disease, whereas EV displays virulence specific to insects (35,36). 70% of residues involved in the SAM binding site and 90% in the GTP binding site are conserved across alphaviruses. Additionally, all common SAM and GTP binding site residues exhibit conservation (Supplementary Figure 1).

The PT compound exhibited the highest binding affinity with a calculated binding energy of - 11 kcal/mol. PT formed hydrogen bonds with Arg70, Arg41, and Asp152, while also engaging in hydrophobic interactions with Ser44, Val243, Tyr285, Phe178, Tyr154, Glu250, His37, Arg92, Gly65, Tyr248 (Figure 1C, 2C) (Table 1). For ABT, the binding energy was slightly lower at -10.3 kcal/mol, primarily establishing a hydrogen bond solely with Arg92 and alongside multiple hydrophobic interactions involving Glu250, Gly65, Ala155, Arg85, Val156, Val153, Tyr154, Ser86, Arg70, His45, Ala69, Lys99, Arg41, His37, Ala40, Phe178, Tyr285, and Tyr248 (Figure 1B, 2B) (Table 1). In contrast, among three, BH displayed the least binding affinity, i.e. -7.2 kcal/mol, with notable interactions observed with Arg41, Asp152, Tyr285, Ser44, and hydrophobic residues including Arg92, Tyr248, His37, and Arg70 (Figure 1C, 2C) (Table 1).

**Figure 1:**
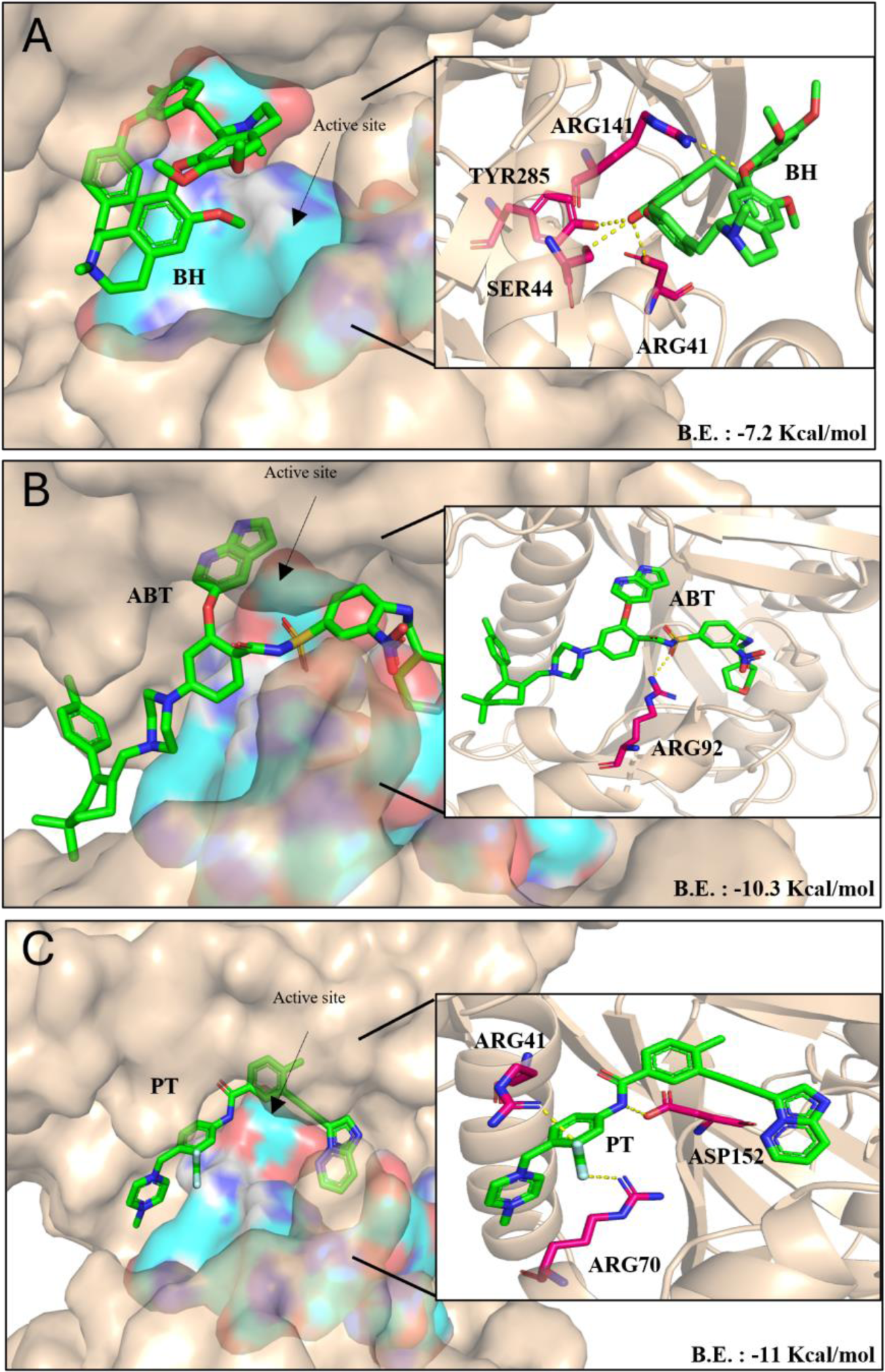
Three-dimensional representation of docked ligands in the enzyme active site (teal surface) of CHIKV nsP1 (wheat surface). (A-C) CHIKV nsP1 interacts with the BH (A), ABT (B) PT (C). Zoomed in figures (A), (B), and (C) show a detailed view of the binding pocket where molecular interactions of ligands (green) with CHIKV nsP1 (wheat coloured protein ribbon/ surface) binding residues (pink colour).

**Figure 2:**
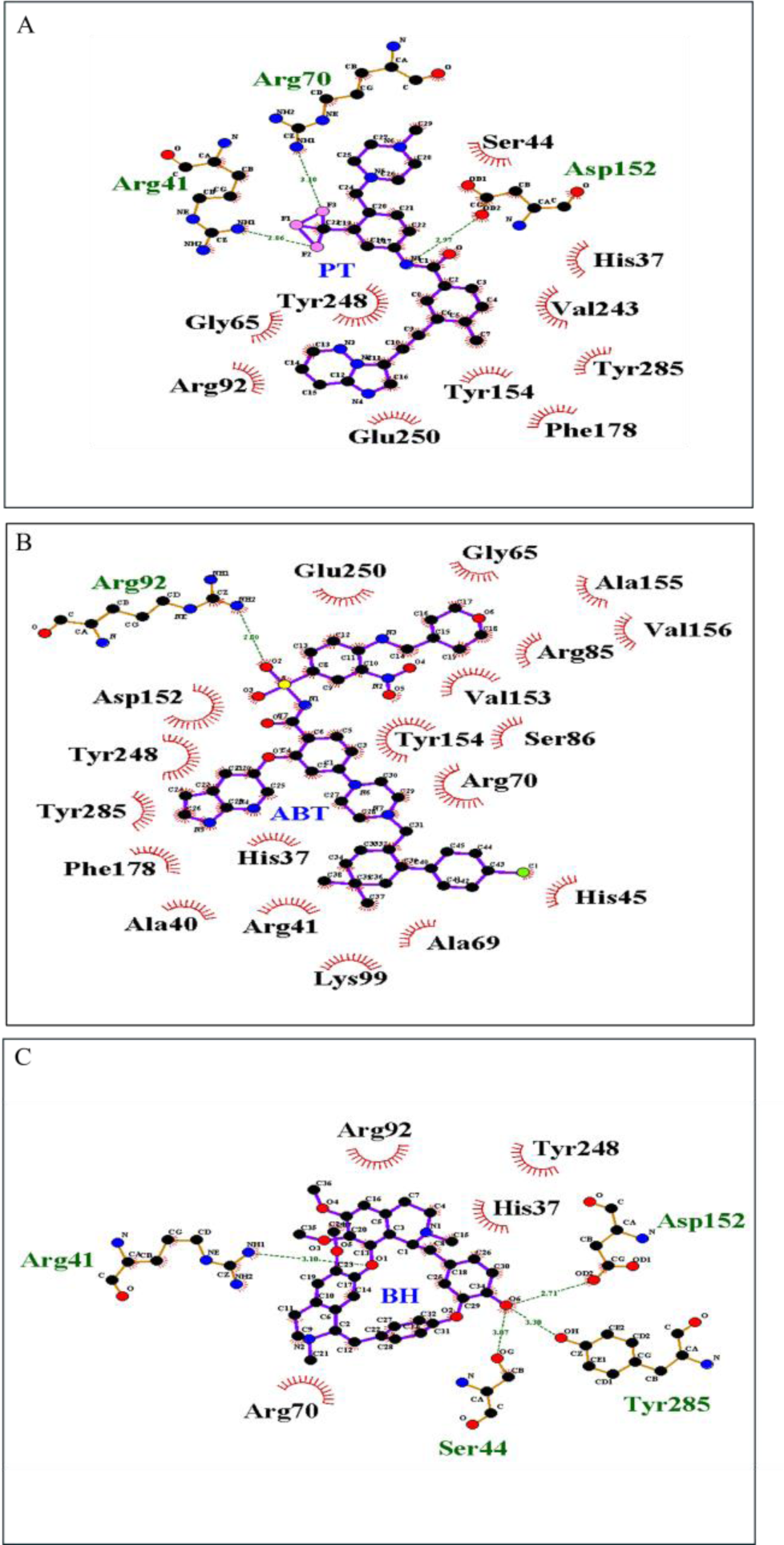
Two-dimensional representation view of docked ligands in the enzyme active site of CHIKV nsP1. CHIKV nsP1 interact with the BH (A), ABT (B) PT (C), H-bonds are shown in green dashed lines with the distance shown in Å. Additional residues forming hydrophobic interactions are shown by brown semicircle with radiating spokes towards the ligands. 2D interaction figures are made using LigPlot+ software.

**Table 1:**
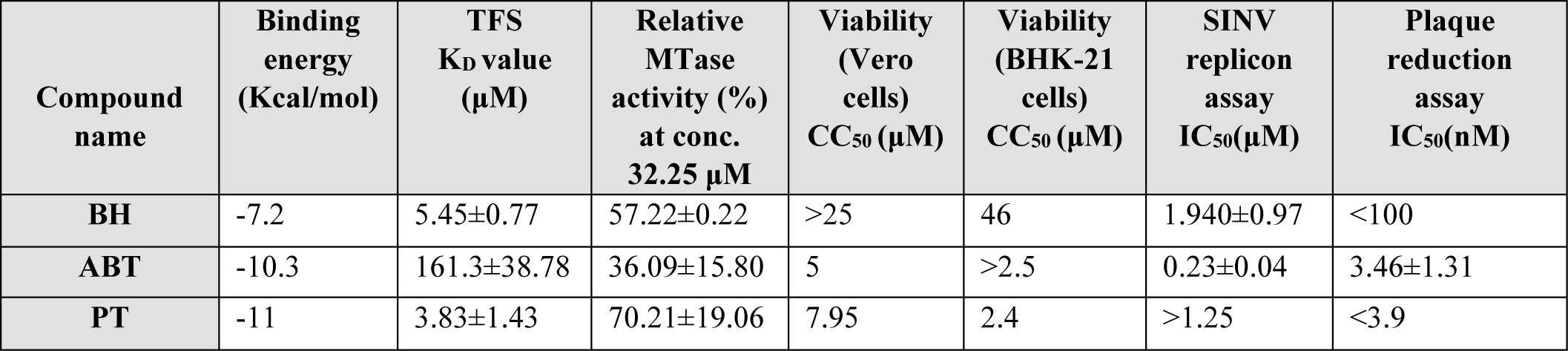
Detailed molecular interactions between the CHIKV nsP1 and ligands.

Molecular dynamics (MD) simulations were employed to study the dynamic stability and conformational changes of CHIKV nsP1 in its apo form and when complexed with ligands. Analysis of the RMSD data demonstrated that equilibrium was achieved within 10 ns for all nsP1-ligand complexes, which then maintained stability throughout the 100 ns simulation period (Supplementary Figure 2). The observed fluctuations ranged from 0.6 to 0.75 nm for the nsP1 apo protein and 0.35 to 0.55 nm for the nsP1-BH, nsP1-ABT, and nsP1-PT complexes. These results demonstrate that ligand binding to the nsP1 protein resulted in stable interactions.

### Binding analysis of molecules to CHIKV nsP1

The interaction of BH, ABT, and PT with purified CHIKV nsP1 was investigated using TFS. The intrinsic fluorescence of the native CHIKV nsP1 proteins was measured utilizing a spectrofluorometer. The binding of these proteins with BH, ABT, and PT at various concentrations was analyzed. Intrinsic fluorescence quenching was observed in increasing compound concentrations (BH, ABT, and PT) in the presence of 1 µM nsP1. In TFS, the red shift indicates a spectral line shifting to longer wavelengths due to increased polarity or decreased hydrophobicity around tryptophan residues. Contrastingly, blue shift signifies a shift to shorter wavelengths due to decreased polarity or increased hydrophobicity. A dose-dependent red shift was observed when BH and ABT interacted with the proteins (Figure 3 D-F). On the other hand, for nsP1 interactions with BH, ABT, and PT, the dissociation constant K_D_ values were ∼5.45±0.77µM, ∼161.3±38.78 µM, and ∼3.83±1.43µM, respectively (Figure 3 A-C).

**Figure 3:**
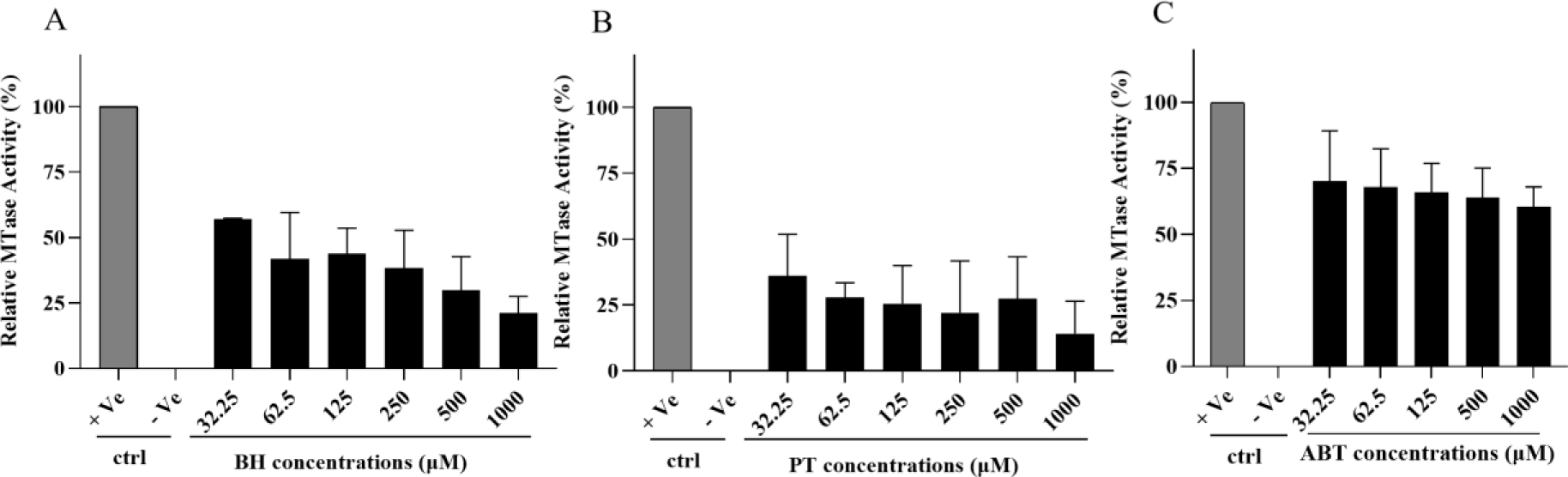
Determination of protein interactions with ligand by TFS. CHIKV nsP1 - (A) BH (B) ABT (C) PT. Intrinsic fluorescence intensity change in protein-compound interactions by TFS. CHIKV nsP1 - (D) BH (E) ABT (F) PT.

### Viral nsP1 MTase enzyme inhibition

The results of the binding study prompted an investigation into the inhibitory effect of these compounds on the MTase activity of purified CHIKV nsP1. A CE-based MTase assay was utilized to quantify SAH levels generated during MTase reactions with BH, ABT, and PT. Relative methyltransferase activity was determined by comparing these levels with a positive control. Additionally, no SAH was detected in the negative control. BH, ABT, and PT dose-dependent decrease confirmed the inhibitory activity of BH, ABT, and PT against a CHIKV nsP1 (Figure 4 A-C). At 32.25 μM concentration, relative MTase activity compared to positive control was ∼57.22±0.22, ∼36.09±15.80, and ∼70.21±19.06 % for BH, ABT, and PT, respectively.

**Figure 4:**
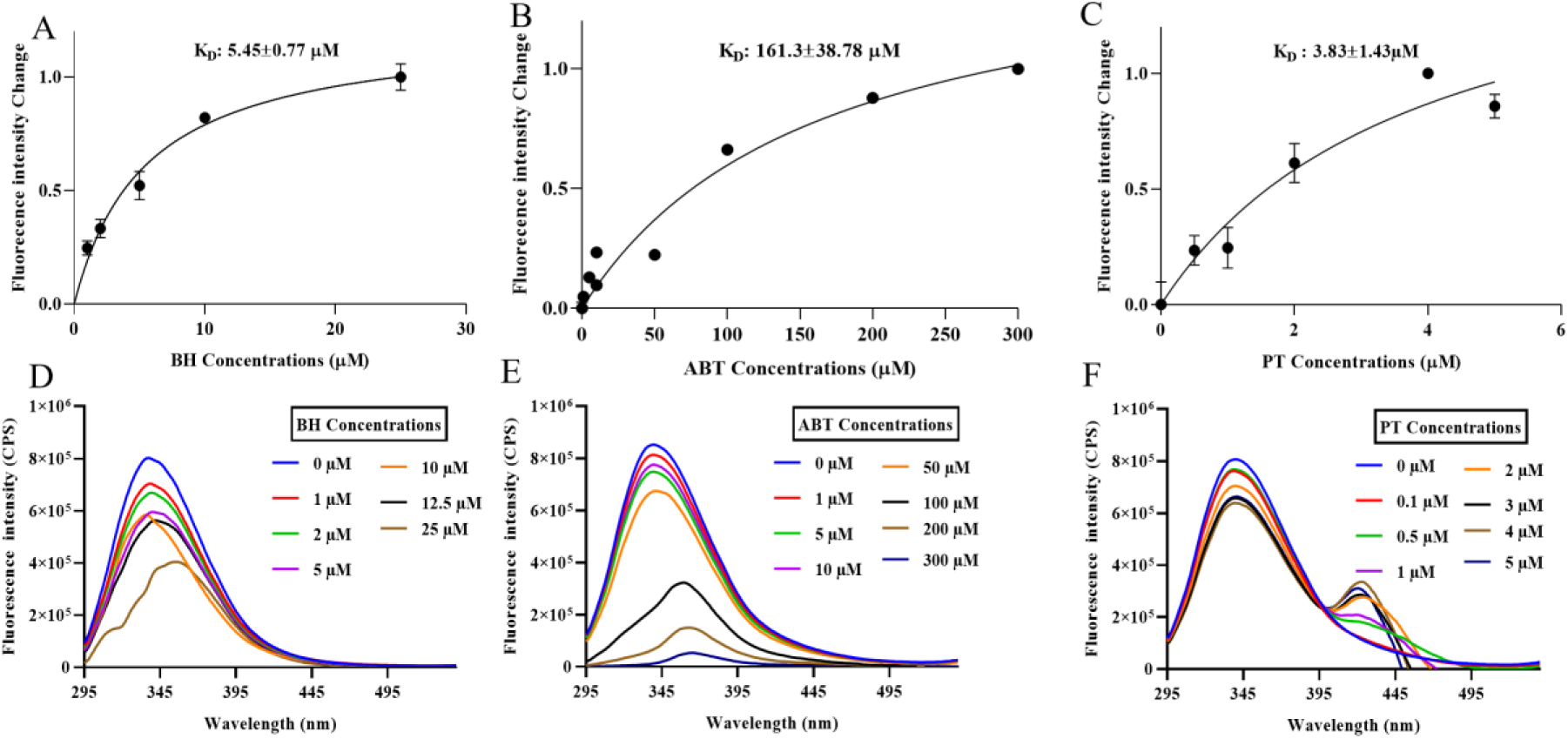
CE-based nsP1 MTase activity inhibition assay. The compounds BH (A), PT (B), and ABT (C) were incubated with 5 µM of CHIKV nsP1 for 1 hour in a reaction mixture containing concentrations ranging from 32.25 µM to 1000 µM. +Ve represents a positive control with no inhibitor added to the reaction. -Ve denotes a negative control where SAM/GTP is absent in the enzyme reaction. Error bars represent the standard error of experimental replicates with n = 3.

**Figure 5:**
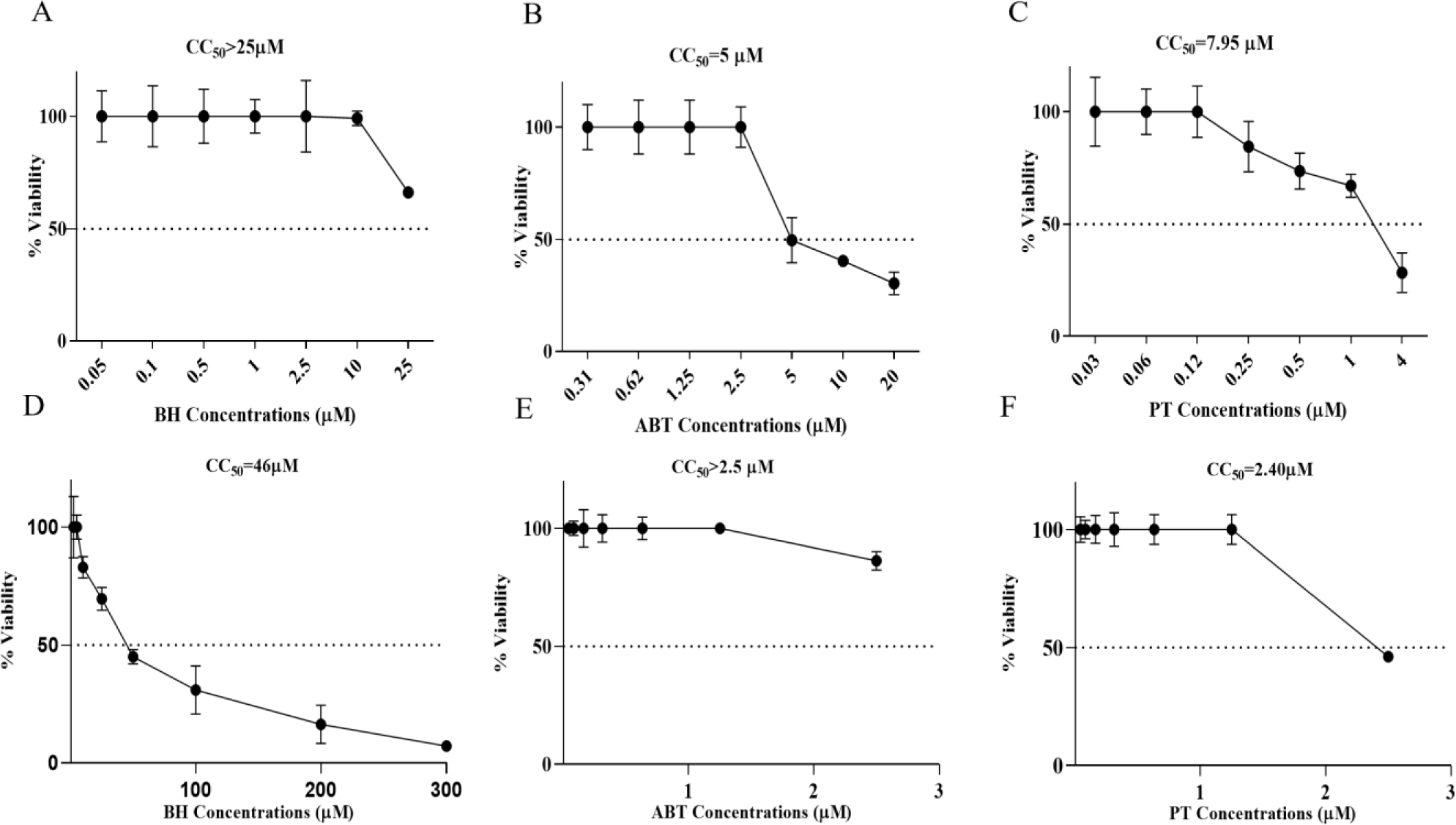
Cell viability analysis by MTT assay (A)BH, (B)ABT and (C)PT depicts the percent cell viability of Vero cells treated compounds for 24 h. (D)BH, (E)ABT and (F)PT depicts the percent cell viability of BHK-21 cells treated compounds for 24 h. MTT assay data represents mean and standard deviations from triplicates (n=3). The compound concentrations showing cell viability more than 90% were selected further to perform antiviral and replicon assays.

### Antiviral efficacy of BH, ABT, and PT against CHIKV

For this study, CHIKV clinical isolate (Accession No. KY057363.1.) was propagated in Vero cells and used in further research (31). Based on the MTT assay, the above 90% viability concentrations of compounds further proceeded for assessment of the antiviral potential of BH, ABT and PT against CHIKV (Figure 6 A-C). A dose-dependent reduction in CHIKV titer was observed in cells treated with BH, ABT, and PT compared to untreated CHIKV-infected cells, i.e., virus control (VC) (Figure 7 A-C). In the case of BH and PT, more than 99 % reduction was seen at 100 and 3.9 nM concentrations, respectively. On the other hand, ABT demonstrated an IC_50_ ∼3.46±1.31 nM obtained from dose-response curve nonlinear regression analysis (Supplementary Figure 2 A-C).

**Figure 6:**
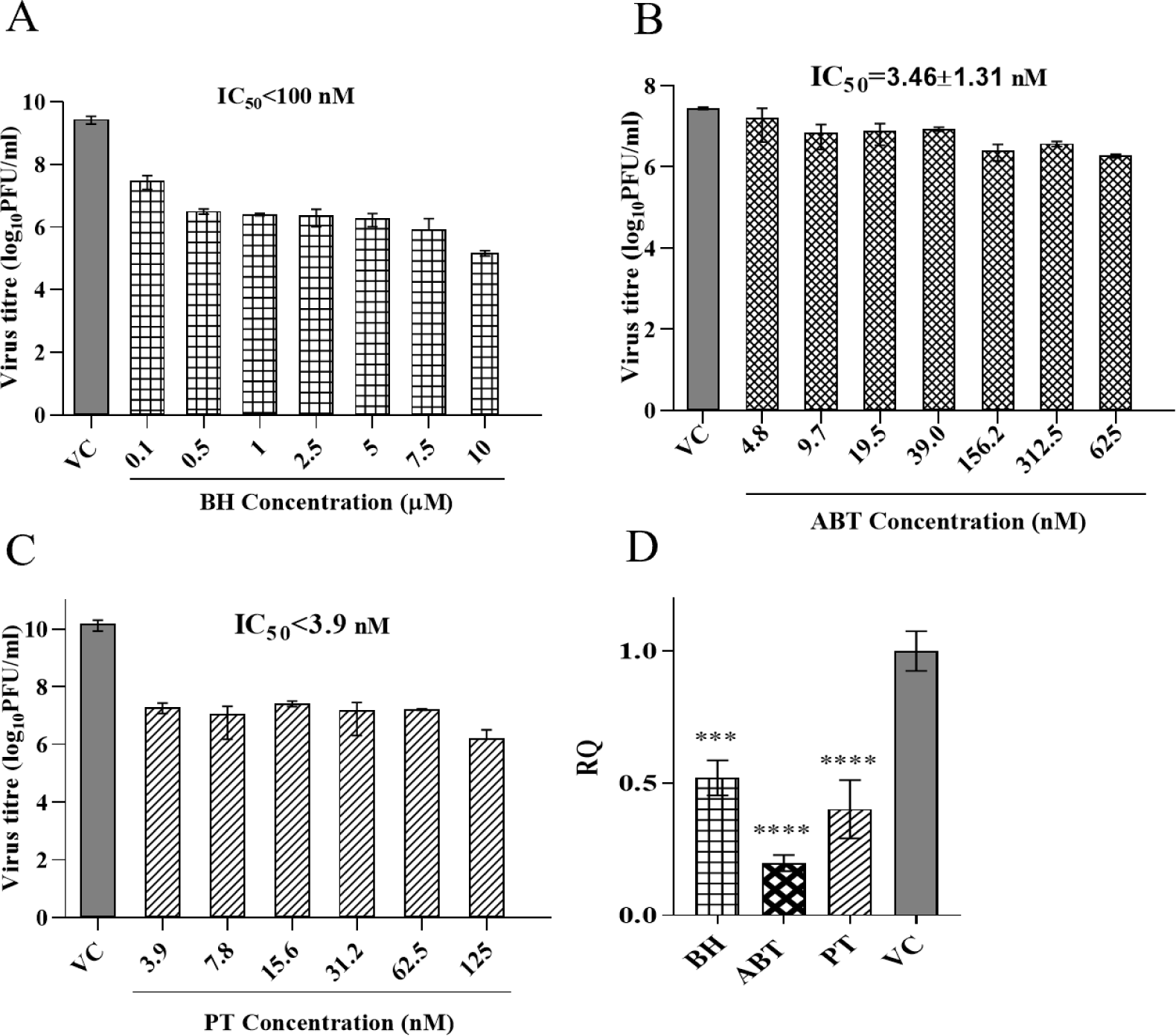
Evaluation of antiviral activity using virus titer reduction profiling of BH (A), ABT(B) and PT(C). Vero cells were treated for 24 h post CHIKV infection, and plaque assay was performed on the collected supernatant. Here, VC represents untreated virus control. (D) qRT-PCR: real-time PCR data shows the viral RNA reduction after 24 h compound (BH 200 µM, ABT 625 nM, and PT 125 nM) treatment compared to virus control in the form of RQ (Relative quantification = 2-ΔΔϹt) value using β-actin as endogenous control. Values are the means, and error bars represent the standard deviation from three independent experiments. Statistical analysis was performed using one-way ANOVA with Dunnett’s post-test., ***P =0.0002, ****P <0.0001.

**Figure 7:**
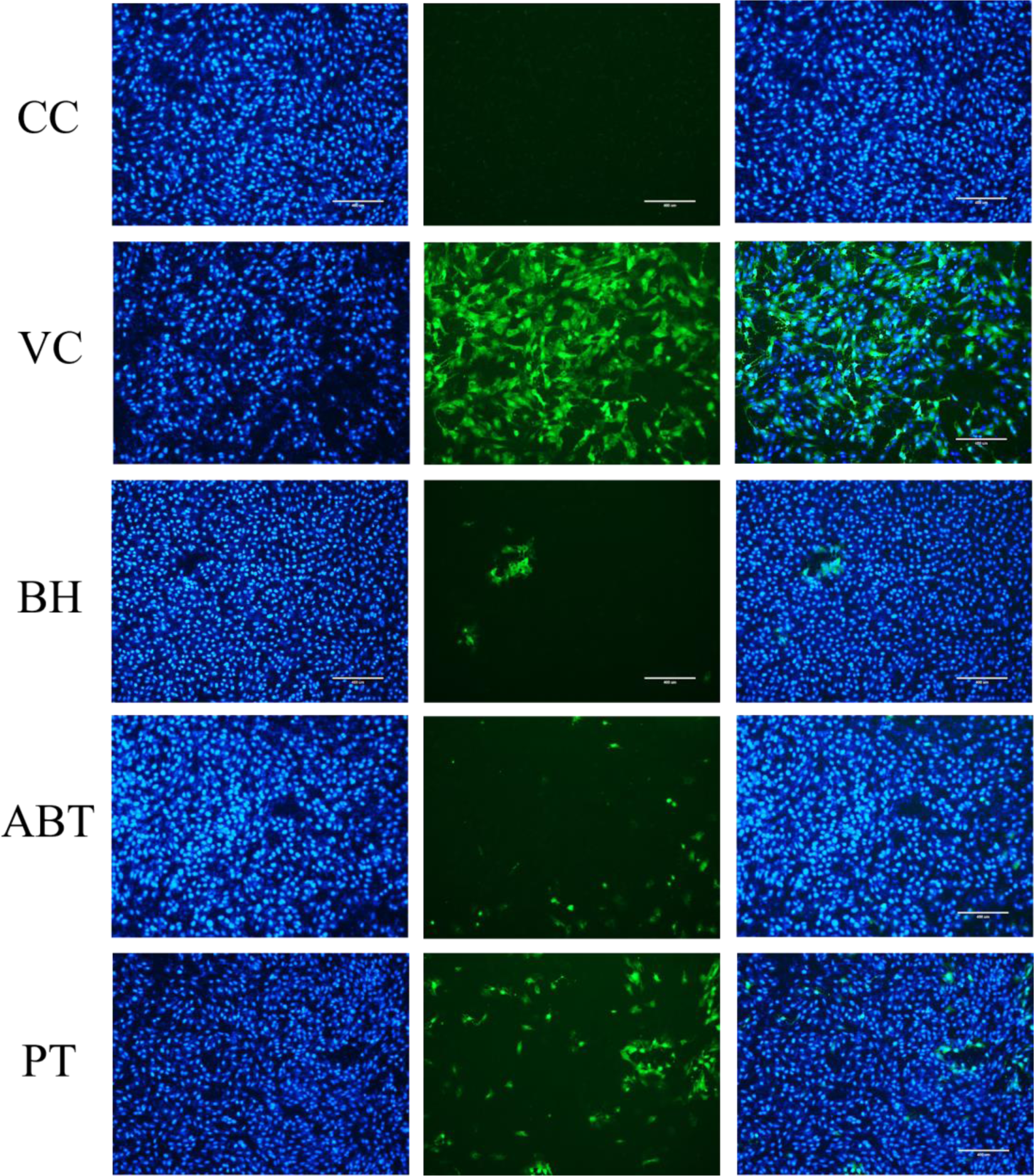
Evaluation of the antiviral effect by immunofluorescence assay (IFA). Immunofluorescent staining of BH (10 µM), ABT (625 nM) and PT (125 nM) for CHIKV. Green fluorescence indicates the virus load, and blue fluorescence indicates the nuclear staining with DAPI with a 10× objective lens. The scale bar is 400 μM.

Here, qRT-PCR was used to validate the antiviral effect of BH, ABT, and PT by quantifying CHIKV RNA levels in the infected cells. qRT-PCR showed a significant reduction in the viral RNA levels when treated with BH, ABT, and PT compounds in an antiviral assay for CHIKV (Figure 7 D). At the mentioned concentrations, BH, ABT, and PT showed a reduction of 2, 5.2, and 2.5 folds in viral RNA, respectively, compared to the VC. IFA analysis was conducted to evaluate the effect of compound treatment on CHIKV-infected cells. Uninfected cells did not exhibit a detectable green fluorescent signal, whereas Vero cells infected with CHIKV (VC) displayed prominent fluorescence. The results showed a decrease in the CHIKV virus titer after treatment with BH, ABT, and PT (Figure 8) compared to VC.

**Figure 8:**
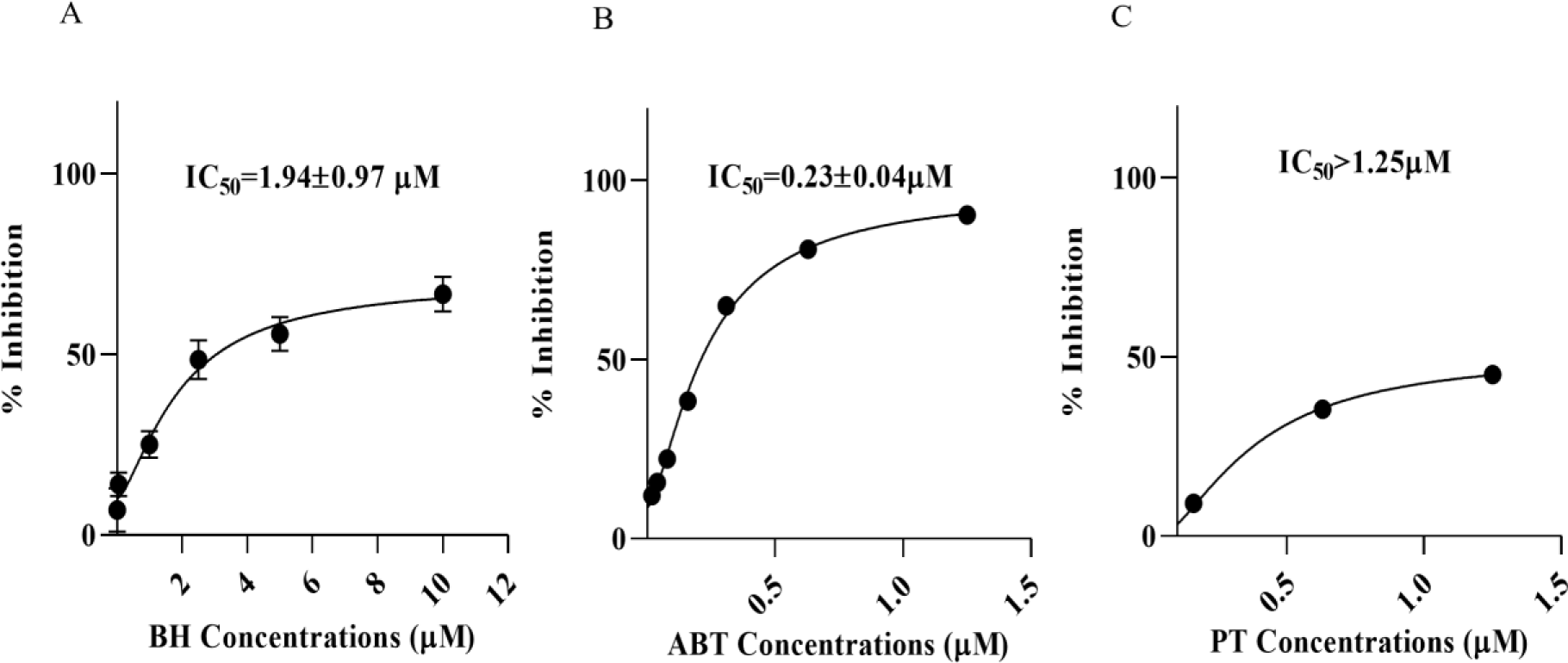
Evaluation of the alphavirus replication inhibitors by SINV replicon assay. (A)BH, (B)PT and (C)ABT. SINV minigenome replicon assay capturing the luciferase activity in lysate derived from BHK-21 cells transfected with capped RNA encoding the SINV-REP reporter replicon. Cells were subjected to 6 h of treatment with inhibitor dilutions. Percentage inhibition values are represented as mean ± SD of three independent experiments. Error bars correspond to the standard deviation from triplicate experiments.

### Inhibition of SINV viral RNA replication by BH, ABT, and PT

Cell viability of BHK-21 in the presence of selected compounds was determined by MTT assay (Figure 6 D-F). Compound concentrations greater than 90% cell viability were employed for the replicon-based luciferase assay. The antiviral activity of compounds was preliminarily evaluated using a SINV-REP system. All three compounds showed a significant decrease in luciferase activity that was used as a surrogate of viral replication in the infected cells. Dose-response curves for IC_50_ values were determined by nonlinear regression fit. The IC_50_ was measured as ∼1.94 ± 0.97 µM and ∼0.23 ± 0.05 µM for BH and ABT, respectively (Figure 9 A,B). The highest average percentage inhibition was measured as ∼45.11±1.97 at 1.25 µM concentration for PT (Figure 9 C). The tested compounds displayed a notable inhibitory effect against the SINV replicon, indicating inhibition of viral RNA genome replication by all the three nsP1 inhibitors.

## Discussion

Arthropod-borne diseases have a significant impact on public health globally. In addition, Chikungunya’s emergence and re-emergence present substantial health risks worldwide (5). Although an FDA-approved vaccine is available against CHIKV, no antiviral therapy has been available. Hence, there is a need to develop antiviral therapy against CHIKV. Extended development timelines and substantial resource allocation hamper the conventional development of antivirals for screening applications (18) The availability of computational tools and structural data enables rapid drug repurposing. This approach is strategically advantageous, leveraging existing medications’ safety profiles and well-characterized pharmacokinetics.

In alphaviruses, nsP1 plays a vital role in viral RNA capping with conserved SAM and GTP binding site (Supplementary Figure 1) (30). The virtual screening of ∼3180 FDA-approved drugs targeting CHIKV nsP1 led to top hits as BH, ABT, and PT could bind in the substrate binding pocket of the nsP1 protein of CHIKV. PT showed the highest binding energy, followed by ABT and BH (Figure 1 A-C). Additionally, all the compounds have formed stable interactions with nsP1 (Figure 2).

TFS was employed to investigated the binding affinities of BH, ABT, and PT to purified CHIKV nsP1. All the compounds have shown fluorescence intensity quenching, indicating conformational changes in CHIKV nsP1 upon interaction. PT has the highest binding affinity for CHIKV nsP1, followed by BH, as evidenced by the K_D_ values (Figure 4 A, C). Among these molecules, ABT has the lowest binding affinity but has shown a sharp red shift in a dose-dependent manner (Figure 4 B, E). Similarly, BH has shown red shift and fluorescence intensity quenching (Figure 4 D). On the other hand, neither red nor blue shift was observed in the case of PT (Figure 4 F). BH, ABT, and PT have shown dose-dependent inhibition in nsP1enzyme activity (Figure 5 A-C). PT has demonstrated the highest inhibition at the lowest tested concentration (32.25 µM), followed by BH and ABT.

This study demonstrated that BH, ABT, and PT have significant anti-CHIKV activity. These compounds showed a reduction in CHIKV infection in the plaque reduction assay (Figure 7 A-C). Moreover, qRT-PCR reveals decreased levels of CHIKV RNA following treatment with BH, ABT, and PT (Figure 7 D). Furthermore, the IFA experiments provided additional validation where BH, ABT, and PT exhibited inhibition compared to the virus control in CHIKV-infected cells (Figure 8).

In the SINV replicon assay, BH, ABT, and PT effectively inhibited viral replication in the compound-treated cells, as evidenced by the decrease in luciferase activity (Figure 9 A-C). These findings suggest that BH, ABT, and PT may function as broad-spectrum inhibitors of alphaviral replication. ABT displayed the most potent antiviral activity among the tested compounds, followed by BH. While PT exhibited a dose-dependent reduction in luciferase activity, its maximum inhibition only reached around 50% at the highest tested concentration. Overall, PT, ABT, and BH have demonstrated strong binding affinities, notable MTase activities, significant antiviral potencies, and replicon inhibition in various assays (Table 2).

**Table 2:**
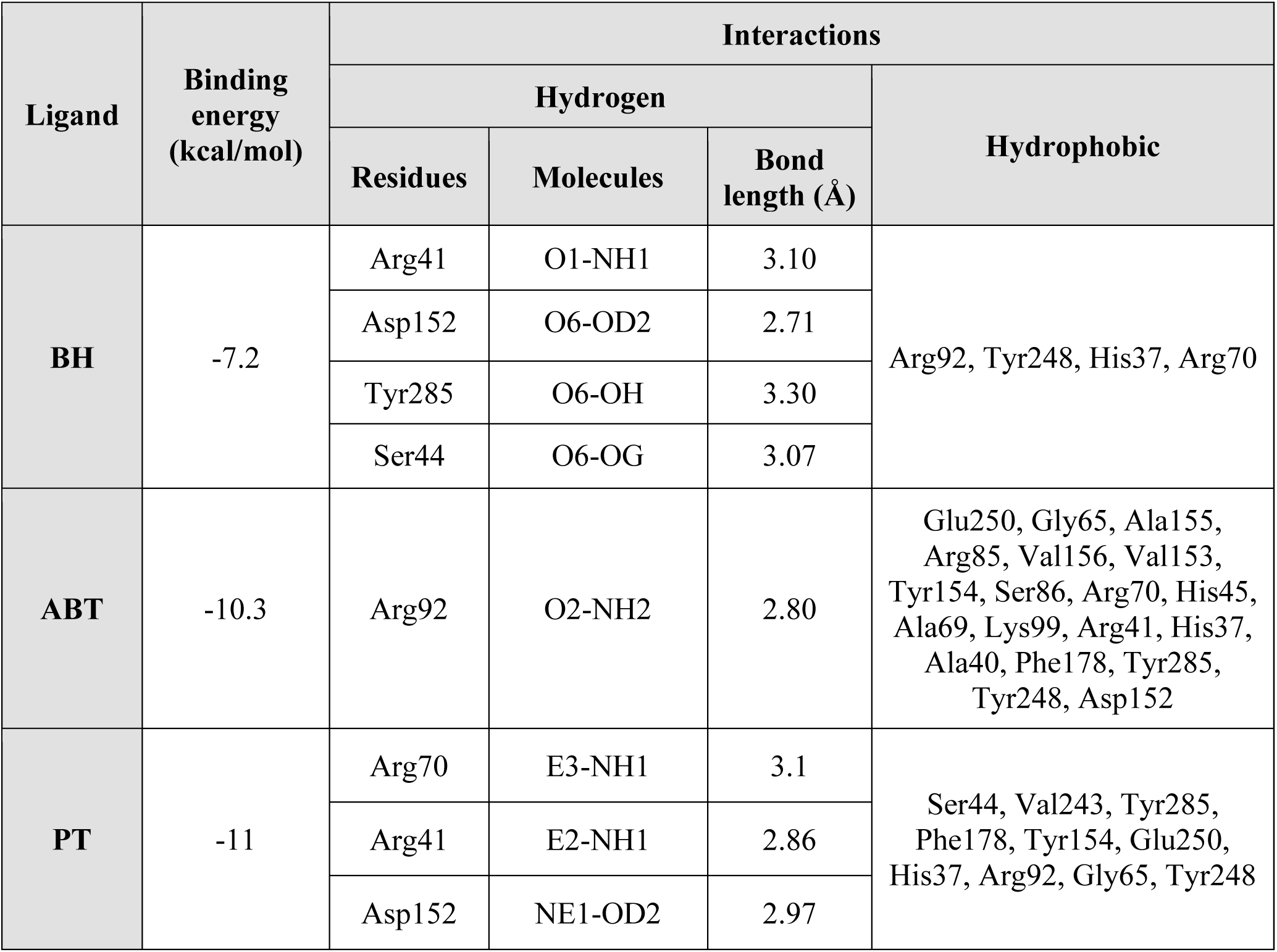
Summary of results from various experiments.

Berbamine is a bis-benzylisoquinoline alkaloid. It has been isolated from the traditional Chinese medicinal herb *Berberis amurensis* (37,38). Additionally, it has been identified in the tree turmeric plant (*Berberis aristata*), which is traditionally utilized in dietary and Ayurvedic practices for its anti-inflammatory and antimicrobial properties. It has demonstrated a cardioprotective mechanism in rodent models (39,40). Various research articles have reported BH as an entry inhibitor in Porcine epidemic diarrhoea virus, Severe acute respiratory syndrome coronavirus 2, Middle east respiratory syndrome coronavirus, Japanese encephalitis virus, African swine fever virus, and Ebola virus (37,41–46). ABT is an FDA-approved drug used primarily for treating acute myeloid leukemia (Supplementary Table 1) (47,48). ABT is a selective inhibitor of the anti-apoptotic protein BCL-2. ABT can specifically induce the premature death of influenza A virus-infected cells (49). PT is a tyrosine kinase inhibitor approved for treating chronic myeloid leukemia and acute lymphoblastic leukemia (Supplementary Table 1) (50). PT can broadly repress latent Human immunodeficiency viruses-1 (HIV-1) reactivation in various cell models by impeding the activation of the AKT-mTOR pathway, which subsequently blocks the interaction between the HIV-1 long terminal repeat and transcription factors (51). These findings suggest that these molecules may target additional host factors involved in CHIKV inhibition, warranting further investigation into their mechanisms of action. However, the inhibition of SINV viral RNA replication in the replicon assay clearly indicates that the viral replication complex, specifically the nsP1 viral enzyme, is the primary target responsible for the observed antiviral activity against alphaviruses. In conclusion, this study underscores the utility of drug repurposing as a promising method for identifying antiviral agents targeting CHIKV viral enzymes.

## Acknowledgments

The authors thank the Department of Biosciences and Bioengineering, IIT Roorkee, for providing central facilities. The authors thank the Macromolecular Crystallographic Facility (MCU) at IIC, Indian Institute of Technology, Roorkee. ST thanks the Department of Biotechnology, Govt of India, for supporting the Bioinformatics Center at IIT Roorkee ref number BT/PR40141/BTIS/137/16/2021. The authors thank the Ashok Soota Molecular Medicine facility, IIT Roorkee. ST also would like to thank DBT, Govt of India “National Network Project Project (BT/PR40142/BTIS/137/72/2023). The authors thank Ms. Shweta Choudhary and Mr. Sanketkumar Nehul for the insightful discussions. The authors would like to acknowledge Ms. Vishakha Singh for her technical assistance in MD simulation. The authors thank the Council of Scientific and Industrial Research (CSIR) and Prime Minister’s Research Fellows (PMRF) scheme for providing financial support. This work was supported by a research grant to ST from ICMR ref no. ISRM/12(46)/2020.

## Supplemental Information

Supplemental information includes three figures, one table

## Authors contribution

Conceptualization, S.T.; methodology, S.T., M.B., and A.S.; experimentation, M.B. and A.S.; formal analysis, S.T.; writing - original draft, M.B. and A.S.; writing - review & editing, S.T.; supervision, S.T.; all authors read, revised, and approved the manuscript.

## Supplementary

**Supplementary Figure 1:**
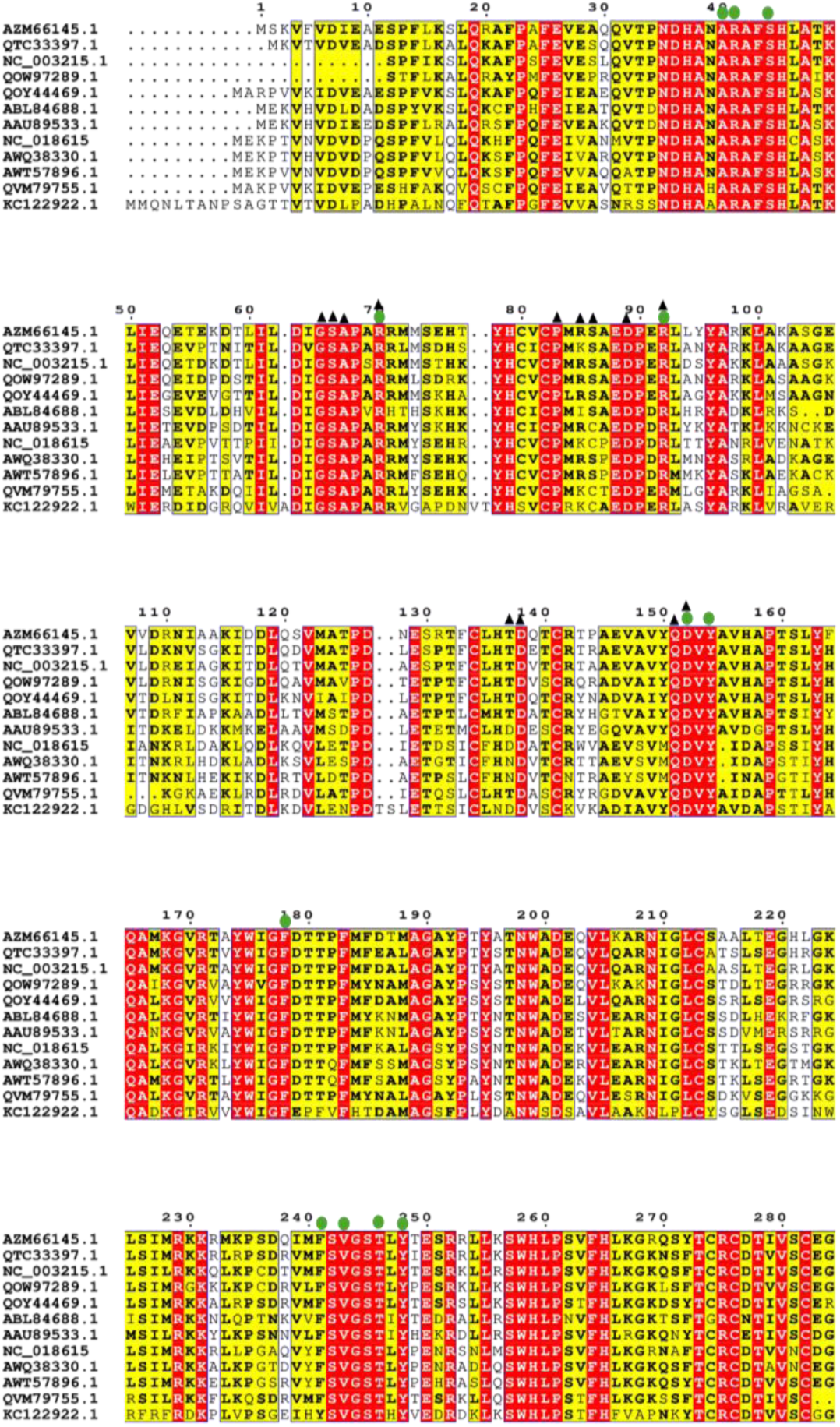
Multiple Sequence alignment of MTase domains of different alphaviruses and flaviviruses. Sequence alignment of nsP1 MTase domain (residues 1 to 285) from CHIKV with the MTase domains of other alphaviruses. MV: AZM66145.1, SFV: NC_003215.1, BFV: QVM79755.1, EV: NC_018615, CHIKV: QOW97289.1, MDV: ABL84688.1, AV: AWQ38330.1, SAV: KC122922.1, RRV: QTC33397.1, VEEV: AAU89533.1, SINV: AWT57896.1, MBV: QOY44469.1. 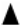 SAM interacting residues 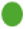GTP interacting residues 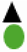 Common residues.

**Supplementary Figure 2:**
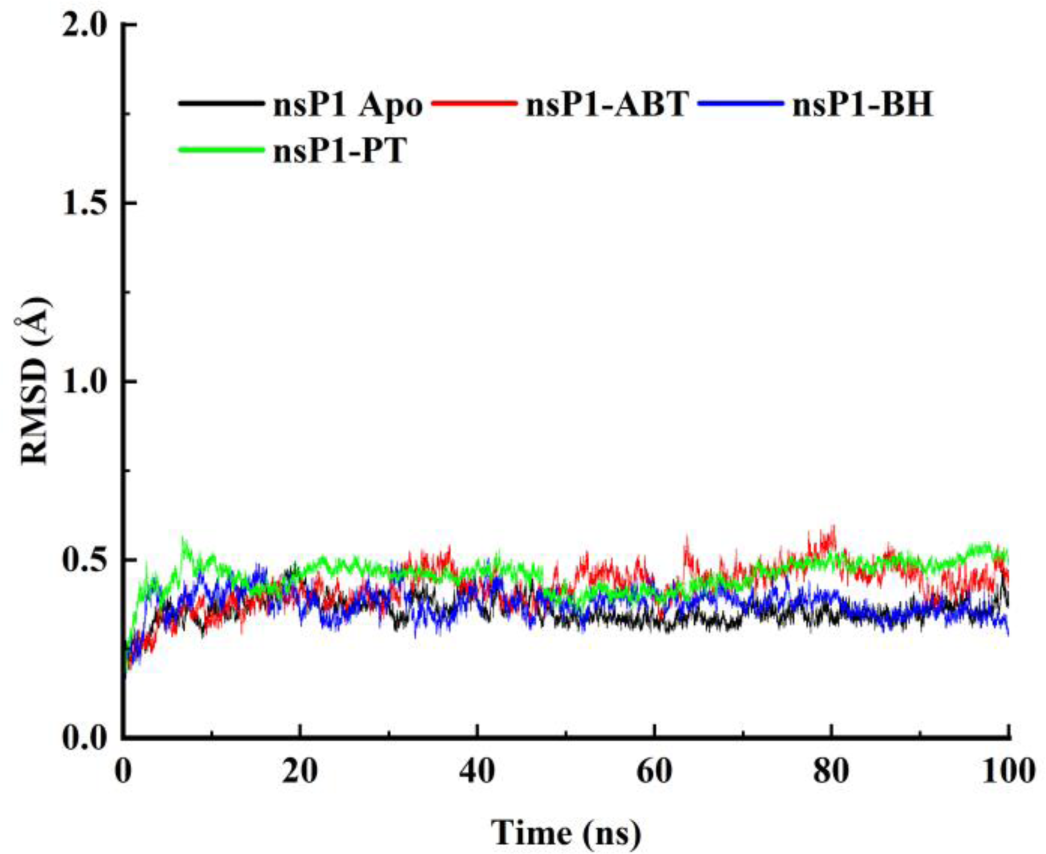
Root Mean Square Deviations (RMSD) graphs of CHIKV nsP1 Apo i.e., native protein, nsP1 - BH or ABT or PT.

**Supplementary Figure 3:**
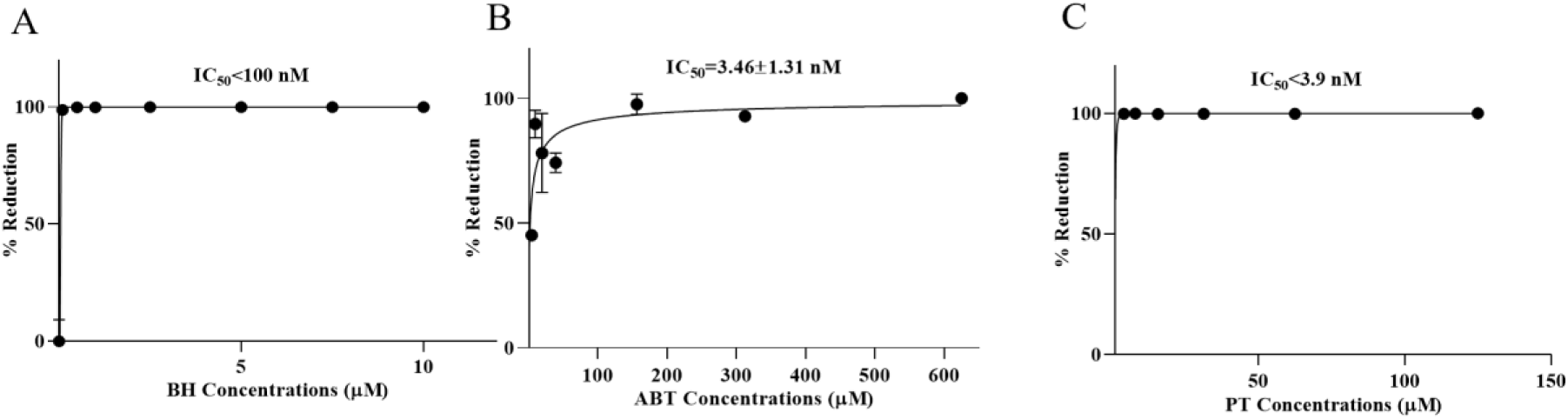
Evaluation of antiviral activity using virus titer reduction profiling of BH (A), ABT(B) and PT(C) using nonlinear regression analysis.

**Supplementary Table 1:**
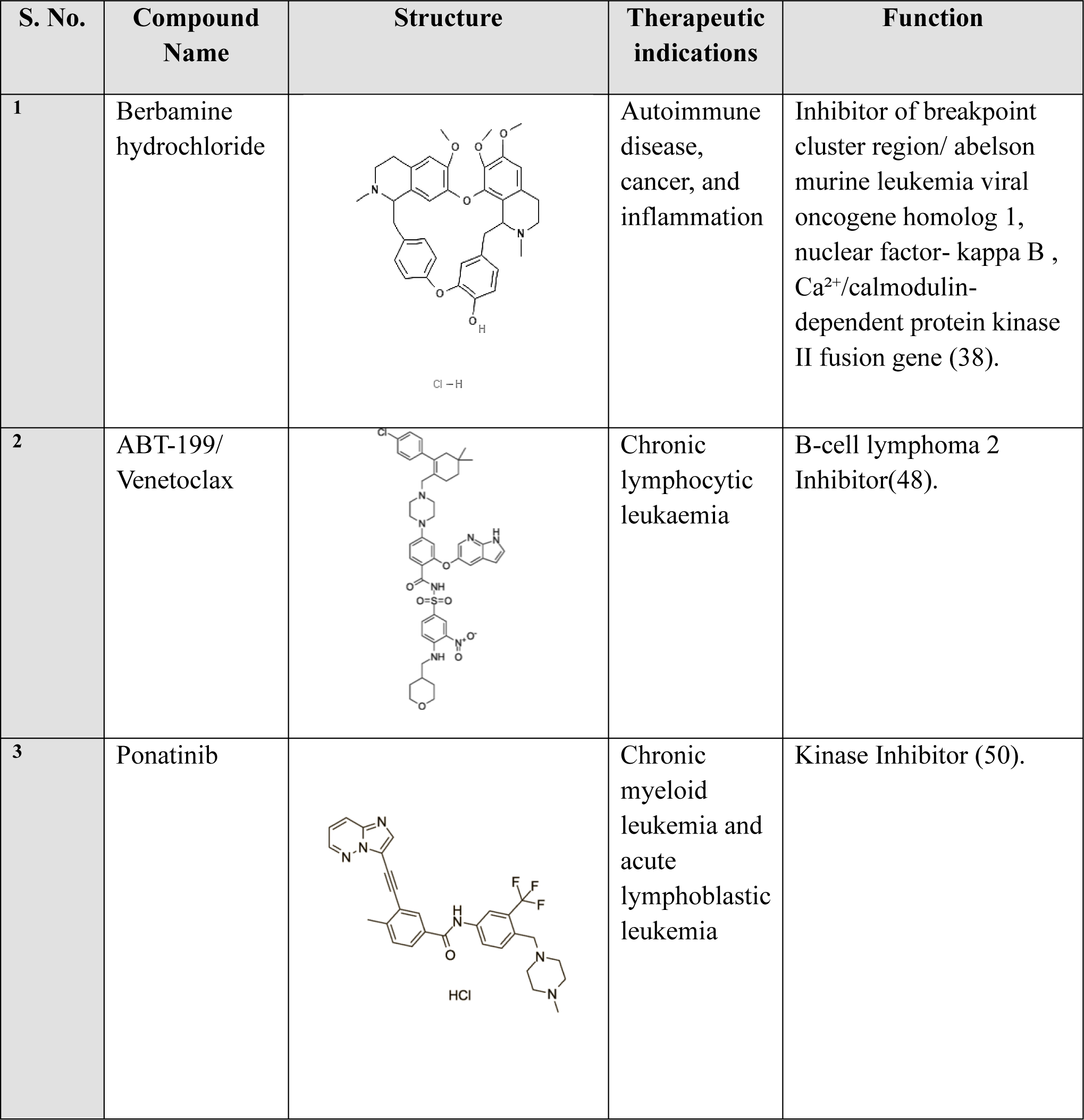
Compounds with Two-Dimensional Structures and Their Therapeutic Indications and Functions.

| S. No. | Compound Name | Structure | Therapeutic indications | Function |
| --- | --- | --- | --- | --- |
| 1 | Berbamine hydrochloride | Cl - H | Autoimmune disease, cancer, and inflammation | Inhibitor of breakpoint cluster region/ abelson murine leukemia viral oncogene homolog 1, nuclear factor- kappa B , Ca<sup>2+</sup>/calmodulin-dependent protein kinase II fusion gene (38). |
| 2 | ABT-199/<br>Venetoclax |  | Chronic lymphocytic leukaemia | B-cell lymphoma 2 Inhibitor(48). |
| 3 | Ponatinib | HCl | Chronic myeloid leukemia and acute lymphoblastic leukemia | Kinase Inhibitor (50). |

